# Long-Term Exposure of Cells to Cdk4 Inhibitor Palbociclib Leads to Chromosomal Aberrations

**DOI:** 10.1101/2023.08.10.552830

**Authors:** Manuel Kaulich, Steven F. Dowdy

## Abstract

Breast cancers are often driven by mutations, alterations and activation of cell cycle regulatory proteins, including the retinoblastoma tumor suppressor protein (Rb), cyclin E and cyclin-dependent kinases (Cdks), especially cyclin D:Cdk4/6 complexes. There are currently three FDA approved Cdk4/6 inhibitors (Cdk4i) for treating breast cancer. The standard treatment protocol is 21 days of continuous Cdk4i treatment, followed by a 7 day cessation period and then repeating the 28 day protocol. We asked the question of what happens to cells that reenter the cell cycle during the 7 day Cdk4i cessation period. Using RPE1 cells containing visual reporter endogenous histone 2B and p27 genes tagged with eGFP and mCherry, we treated the cells with a Cdk4i, Palbociclib for 1 to 42 days that spanned the clinical exposure, followed by drug release (washout) and video microscopic analysis. Surprisingly, we found that as little as 4 days of Cdk4i treatment and release resulted in a significant increase in micronuclei and multinucleated cells that had reentered the cell cycle. The peak chromosomal aberration occurred between 14 and 35 days, a timing that spans the clinical dosing regimen.These observations raise questions concerning the potential that cycling patients on and off of Cdk4 inhibitors may generate gross chromosomal changes to tumor cells that reenter the cell cycle during the 7 day clinical cessation (washout) period and thereby increase the potential to initiate secondary oncogenic events.

## Introduction

For women in the US, breast cancer remains the most common form of cancer and the second most common cause of death by cancer [1]. Breast cancers are often driven by mutations, alterations and activation of cell cycle regulatory proteins, including the retinoblastoma tumor suppressor protein (Rb), cyclin E and cyclin-dependent kinases (Cdks) [2,3]. Previously, Cyclin D:Cdk4 complexes were thought to inactivate Rb by hypo-phosphorylation [4,5]; however, recent evidence from our lab and others have now shown that Cyclin D:Cdk4 only quantitatively mono-phosphorylate Rb in early G1 phase of the cell cycle and that the 14 mono-phosphorylated Rb isoforms each selectively bind cellular targets [6–8]. In contrast, activation of cyclin E:Cdk2 complexes at the Restriction point perform the initial Rb inactivation by hyper-phosphorylation, triggering release of E2F transcription factors, progression into late G1 phase and subsequent entry into S phase [6,7]. While it remains unclear what the non-Rb targets of Cyclin D:Cdk4 are during early G1 phase to drive cell cycle progression, it is clear that continuous Cyclin D:Cdk4 activity is a required driver of early G1 cell cycle progression in many cancers, including breast cancer [9].

The FDA approved the first Cdk4 inhibitor (Cdk4i), Palbociclib (Ibrance), as a breakthrough therapeutic in 2015 to treat estrogen receptor positive (ER+) breast cancer [10]. This was followed up in 2017 with FDA approval for Cdk4i use in ER+/HER2-negative breast cancer [11]. Given the importance of cyclin D:Cdk4 in driving cancer, it is not surprising that there are now three FDA approved inhibitors [12]. The standard Cdk4i treatment is 21 consecutive days in combination with anti-estrogen drugs, followed by a 7 day cessation, and then repeating the 28 day protocol [13]. Previously, palbociclib was shown to be clastogenic in an in vitro micronucleus assay in Chinese Hamster Ovary (CHO) cells as well as in an in vivo in the bone marrow of male rats dosed at ≥100 mg/kg/day for three weeks [14]. However, it was also reported to not be mutagenic in an in vitro bacterial reverse mutation (Ames) assay and also did not induce chromosomal aberrations in an in vitro human lymphocyte chromosome aberration assay [14,15].

During use for cell cycle experiments in the laboratory, Cdk4i are typically added for only one or two days [6]. Based on pilot observations in our lab, we investigated the consequences of treating cells with longer durations of Cdk4i (3 to 42 days) that spanned the 21 day clinical treatment protocol, followed by release from the drug, an approach that we have not seen previously published. We found that cells reentering the cell cycle after Cdk4i for as short as 4 days resulted in a high propensity to generate severe mitotic defects, leading to micronuclei and multinucleation. These observations raise questions concerning the potential that cycling patients on and off of Cdk4 inhibitors may generate gross chromosomal changes to tumor cells that reenter the cell cycle during the 7 day clinical cessation (washout) period and thereby increase the potential to initiate secondary oncogenic events.

## Materials and Methods

### Cell Culture and synchronization

Human hTERT-RPE1 cells were cultured at 37°C, in 5% CO2 atmosphere and DMEM/F12 media (Life Technologies), supplemented with 10% heat-inactivated fetal bovine serum (FBS) and penicillin/streptomycin (100 μg/ml). For synchronizations, cells were treated with 2 mM thymidine (Sigma-Aldrich), 4 mM hydroxyurea (Sigma-Aldrich), 9 uM RO3306 (Sigma-Aldrich) for 24 hr prior to release, or with 1 μM Palbociclib (PD-0332991, IBRANCE®; Selleck Chemicals) for 48 hr if not indicated otherwise. Cells were released by washing four times with full media and subsequently analyzed or treated as indicated.

### Time-lapse video microscopy, data analysis

For cell cycle analysis, hTERT-RPE1 cells were seeded into either 6-well or 96-well chambers, treated as indicated and imaged using a Nikon ECLIPSE Ti microscope equipped with a CoolLED pE-1 excitation system and a ×20/0.75 air Plan Apo objective (Nikon). For single cell tracing, hTERT-RPE1 cells were seeded into 96-well chambers, treated as indicated and imaged using the high-throughput imaging systems Yokogawa CQ1 or Yokogawa CV7000S (Yokogawa Electric Corporation). Images were acquired by simultaneously recording of 2×3 or 3×3 positions every 10 or 20 min and automatically stitched and processed by Fiji (version 2.0.0-rc-43/1.50e) [16]. Each experiment was carried out in biological replicates (triplicates) on independent days. Each replicate recorded 6 stills of a single well as part of a 96 well plate that were stitched together using CellPathfinder to analyze a minimum of 100 cells per well, with 2 wells per replicate. We have added this information to the experimental design section. To figure 2a, nuclear H2B-eGFP signal intensities were normalized to the mean nuclear H2B signal intensity of time point day 1. To figure 2c and 2d, total values of nuclear p27-mCherry signal intensities were normalized to the mean nuclear p27-Cherry signal intensity of time point day 1. To figure 2e, nuclear size was normalized to the average nuclear size of cells of time point day 3.

### Single cell tracing

Images were imported to Fiji and stack converted to binary files with the ‘Li’ method and ‘Dark’ background and through calculation of thresholds for each image. Subsequently, the TrackMate plugin (v2.8.1) [17] was used for single cell tracing with the following settings: DoG detector, 20 um blob diameter with a threshold of 0.5, 15 μm of linking distance, 15 μm of gap-closing maximal distance and 2 frames for maximal gap-closing.Single cell traces were exported as csv-files for further analysis.

### Bioinformatics and biostatistics

Movies from microscope were processed and exported as .xls files for H2B measures. Files were merged by using a 1100 pixel radius around every H2B data point for every cell. All cells without time point zero were filtered out, to avoid complicated alignment algorithms. To normalize for measurement inaccuracy and to fill missing data points each data point for H2B was smoothed by the median within a +/- 5 time point windows. Next, all cells with a measured time of less than 10 hr (60 time points) and all cells with no H2B-GFP signal were filtered out. Finally the distribution of H2B intensities over time per cell was scaled down to values between 0 and 1. All plots show the data smoothed with the median and normalized to 1. To show graphs about trends in the population, all H2B values for all cells at every specific time point were averaged, using the mean. For defining a cell as cycling an increase of 60% between the lowest H2B and the highest H2B value was required. Cells with no H2B signal increase or less than 60% were defined as not cycling. To calculate the turning point for the H2B signal the global minima was calculated, requiring the last 10 time points to decrease and the next 10 time points to decrease in signal. Nuclear size was determined based on the H2B-GFP reporter protein area of the nucleus using Fiji (version 2.0.0-rc-43/1.50e) [16].

## Results

### Tagging endogenous Histone 2B and p27 as single cell reporter for cell cycle entry

The use of fluorescent cell cycle reporter systems has greatly enhanced our understanding of cell cycle regulation [18,19]. However, overexpressed reporter transgenes can biologically interfere with the very processes that they are intended to visualize. In contrast, endogenous proteins escape this obstacle, but are difficult to study on the single cell level. To investigate the long-term exposure risks that cells experience during Cdk4i treatment and after release back into the cell cycle and visualize cell cycle progression (and DNA content) in an unperturbed manner, we first generated a microscopic visual reporter cell line to allow for single cell analysis. Histone transcription and translation is tightly coupled to DNA replication [20]. Therefore, we knocked-in eGFP into the translational termination site of the endogenous Histone 2B (H2B) genomic locus (Sup Figure S1). RPE1 H2B-GFP (clone #4) cells divided, excited and entered the cell cycle on par with parental RPE1 cells (Sup Figure S1). H2B-GFP nuclear content mirrored DNA content as determined by propidium iodide staining during the cell cycle and in response to exposure of cell cycle arrest agents (Figure 1; Sup Figure S2). To this cell line, we also knocked-in mCherry into the translational termination site of the p27 Cdk inhibitor CDKN1B genomic locus [21,22]. p27 is a nuclear protein in dividing cells; however, we and others have previously shown nuclear export of p27 into the cytoplasm and can thereby serve as an excellent reporter of cellular phenotypes [23-26].

**Figure 1.**
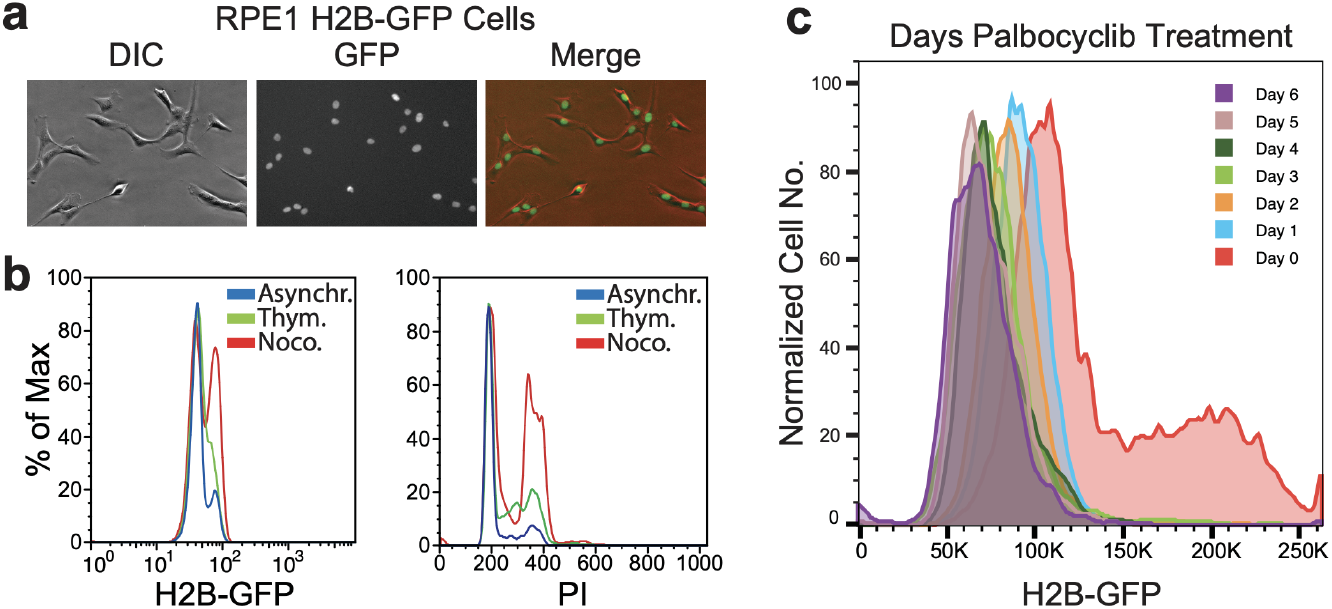
hTERT RPE H2B-GFP Cells Used for Single Cell Analysis. (**a**) Microscopy of H2B-GFP reveals exclusive nuclear staining. (**b**) H2B-eGFP cells were synchronized with Thymidine or Nocodazole and subsequently analyzed by FACS and PI staining. Note: the GFP signal correlates with the cell cycle.

### Cellular consequences of long-term Cdk4 inhibitor treatment

To investigate the potential cellular consequences of long-term exposure to a Cdk4i, we treated RPE1 H2B-eGFP/p27-mCherry reporter cells with 1 μM Palbociclib for 1 to 6 days and 7 to 42 days. 1 μM Palbociclib is the standard concentration used to arrest cycling cells [6]. Treated cells of >4 days showed an induced cell flattening phenotype with increased cellular area (Figure 2A), previously associated with a senescence-like morphology [27]. As expected, the nuclear H2B-eGFP amount remained constant during the entirety of Palbociclib exposure from 7 to 42 days (Figure 2B). However, starting at day 4 of treatment, we also observed a dramatic increase in nuclear size (measured by nuclear H2B-eGFP signal) (Figure 2E). By day 5 of Palbociclib treatment, the cells had essentially doubled their nuclear size (Figure 2E). Since nuclear H2B-eGFP levels did not change due to Palbociclib treatment (Figure 2B), H2B-eGFP served as an internal control for nuclear and cytoplasmic p27-mCherry signals. After an initial gain in nuclear p27-mCherry signal (from days 2-5), we then observed a net loss of nuclear p27-mCherry and an increase in cytoplasmic p27-mCherry signal (Figure 2C,D). Maximal cytoplasmic p27-mCherry signal was reached after 28 days and did not change thereafter (Figure 2D). Thus, long-term Palbociclib exposure induced a nuclear to cytoplasmic translocation of p27-mCherry and cytoplasmic p27 correlates with poor patient prognosis in multiple malignancies [28–31].

**Figure 2.**
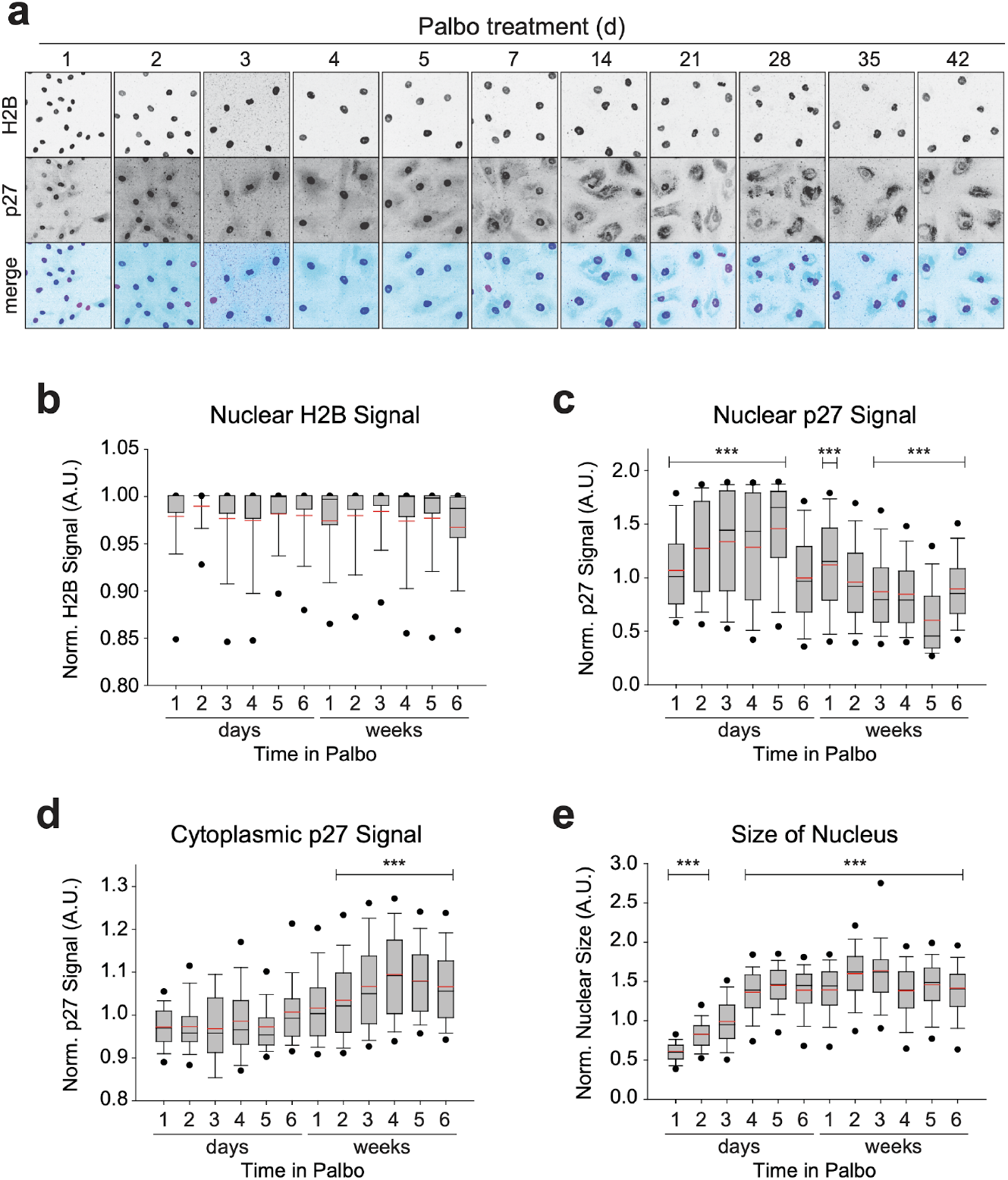
Nuclear growth as stationary consequences of long-term Palbociclib exposure. (**a**) H2B-eGFP and p27-mCherry signal of cells treated with 1 μM Palbociclib for the indicated period of time. (**b**) Nuclear H2B-eGFP does not change its localization and intensity in a Palbociclib-dependent manner. (**c**) Nuclear p27-mCherry increases in intensity for the first 4 to 5 days of Palbociclib treatment before a sharp decline. (**d**) Cytoplasmic p27-mCherry signal rises in intensity after 1 week of continuous Palbociclib treatment. (**e**) Normalized nuclear size as measured by H2B-eGFP signal of single cells. Asterisks (based Mann-Whitney-U-Test): ***: p ≤0.001. Mean (black line) and average (red line).

Palbociclib is an FDA approved Cdk4i for the treatment of postmenopausal breast cancer requiring 21 consecutive days of treatment, followed by a 7 day cessation period and then repeating the 28 day protocol. Therefore, we investigated the potential consequences of long-term Palbociclib treatment and release (cessation) on cells. To do so, we treated RPE1 H2B-eGFP/p27-mCherry reporter cells with Palbociclib for a duration that spanned the 21 day clinical treatment protocol and then released (wash-out) them from Palbociclib arrest and monitoring H2B-eGFP and p27-mCherry signals over the next 48 hr. As we have previously observed, cells treated with 1 μM Palbociclib for 1 or 2 days and released showed unperturbed cell cycle kinetics and completed their first and second cellular divisions after 24 hr and 47 hr of Palbociclib wash out, respectively (Figure 3A). However, cells treated for 3 to 6 days with Palbociclib showed both a delayed entry into their first round of division and were unable to complete it by the time we stopped recording at 48 hr (Figure 3A). Consistent with this trend, cells treated with Palbociclib for 7 to 42 days and released by wash-out displayed a stronger time-dependent decrease in the total number of cells able to re-enter and complete cellular division (Figure 3B).

**Figure 3.**
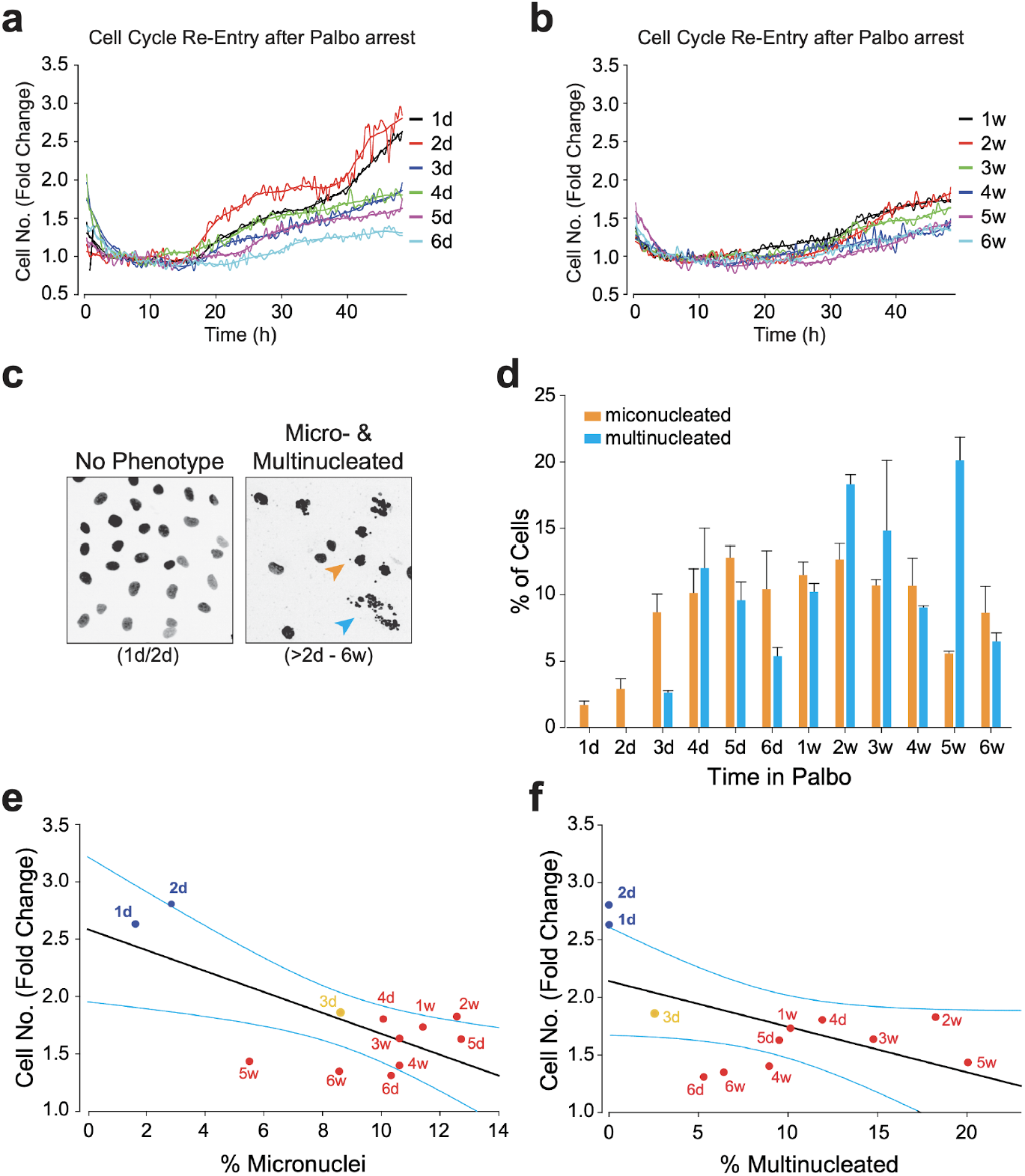
Long-term Palbociclib exposure interferes with cell cycle re-entry and results in mitotic errors. (**a**,**b**) Cells were continuously exposed to 1 μM Palbociclib for up to 6 days (a) or 6 weeks (b) and analyzed for their ability to re-enter the cell cycle and undergo mitotic divisions. (**c**,**d**) Fixed stills from representative movies having micro- and/or multinucleated cells after long-term Palbociclib exposure and their occurrence after Palbociclib release and mitotic division. Orange: micronuclei, blue: multi-nucleation. Bars of (d) represent standard deviation. (**e**,**f**) Correlation between percentage of phenotype and the cells ability to divide (fold change). Blue lines: 95% confidence interval.

While cells that did not enter the cell cycle after Palbociclib release displayed no overt microscopically observable phenotypes, cells that did re-enter the cell cycle displayed severe mitotic phenotypes. We were able to score for both micronuclei and multinucleated phenotypes as a consequence of improper chromosome separation during mitosis (Figure 3C). Interestingly, both phenotypes reached a maximum of ∼10-15% of the population after 4 to 5 days of Palbociclib treatment (Figure 3D) and strongly correlated with an individual cell’s ability to re-enter the cell cycle (Figure 3E,F). These data demonstrated that long-term Palbociclib treatment of immortalized, but non-tumorigenic, human hTERT-RPE1 cells strongly interfered with their ability to re-enter the cell cycle and that cells capable of re-entering the cell cycle after Palbociclib release were prone to chromosome segregation errors during mitosis.

## Discussion

Clinical treatment of breast cancer patients with the Cdk4 inhibitor Palbociclib is a 21 day duration, followed by a 7 day cessation (or release) of treatment, and then repeating the 28 day protocol [10–13]. Our data point to a sharp time window during which cells arrested for 2 days or less were able to efficiently reenter the cell cycle free from gross chromosomal alterations. However, we identified 4 days of Cdk4i Palbociclib treatment and release as the beginning of a critical time point after which cells showed a strong decrease in both their ability to reenter the cell cycle and those cells that eventually pass the early to late G1 restriction point displayed severe mitotic defects, leading to micronuclei and multi-nucleation. This is in line with previous findings showing that Cdk4/6 inhibition has been associated with permanent growth arrest and senescence in some tumor cell types [23,26,28-30]. Interestingly, within this melanoma model, senescence-associated marks (SA-βgal, SASP, SAHF, and ATR-X foci formation) increased at day 4 of Cdk4i Palbociclib treatment, coinciding with our finding that after 3 days of Cdk4i exposure, cells lose their ability to re-enter the cell cycle [26]. In line with this, RPE1 cells exposed to Palbociclib for more than 3 days lost their early G1 phase ‘memory’ and started to demonstrate morphological signs of senescence, such as a flatten cell phenotype. Accordingly, we noticed a significant increase in nuclear size that did not correlate with an increased PI staining, indicating a stable early G1 phase arrest. Thus, increasing nuclear size scores with both the inability to reenter the cell cycle and the appearance of micronuclei. While p27 translocation out of the cytoplasm appears to lag nuclear size increase, we cannot rule out that there is a critical threshold level of p27 in the nucleus that once below, triggers the generation of micronuclei. However, the molecular link to Cdk4i Palbociclib arrest/release and our identified mitotic defects awaits further analysis. Lastly, because we focused our efforts on the Palbociclib Cdk4i, we cannot rule out at this time whether these effects are due specifically to Palboiclib or are class-wide effects of all Cdk4 inhibitors, though we suspect that will be the case. Taken together, our observations raise questions concerning cycling patients on and off of the Palbociclib Cdk4 inhibitor during the standard protocol and thereby increase the potential for clastogenic events during the subsequent passage through M phase, leading to micronuclei and multinucleation. While some or many of these cells may be removed from the body by senescence or other processes, micronuclei are known to undergo chromothripsis, a potent cancer driving event [32].

## Supplementary Materials

The following are available online at www.mdpi.com/xxx/s1, Figure S1: hTERT-RPE1 H2B-eGFP cells, Figure S2: p27-mCherry hTERT-RPE1 Cells.

## Author Contributions

Conceptualization, M.K. and S.F.D.; formal analysis, M.K.; writing— original draft preparation, M.K and S.F.D..; writing—review and editing; funding acquisition, S.F.D. All authors have read and agreed to the published version of the manuscript.

## Funding

This research was funded by Pfizer-CTI.

## Acknowledgements

We are particularly thankful to D. Jenkins, A. Desai and A. Shiau of the San Diego Ludwig Institute for their help with microscopy and the Developmental Studies Hybridoma Bank, University of Iowa, for antibodies.

## Conflicts of Interest

The authors declare no conflict of interest.

## Figure legends

**Supplemental Figure S1.**
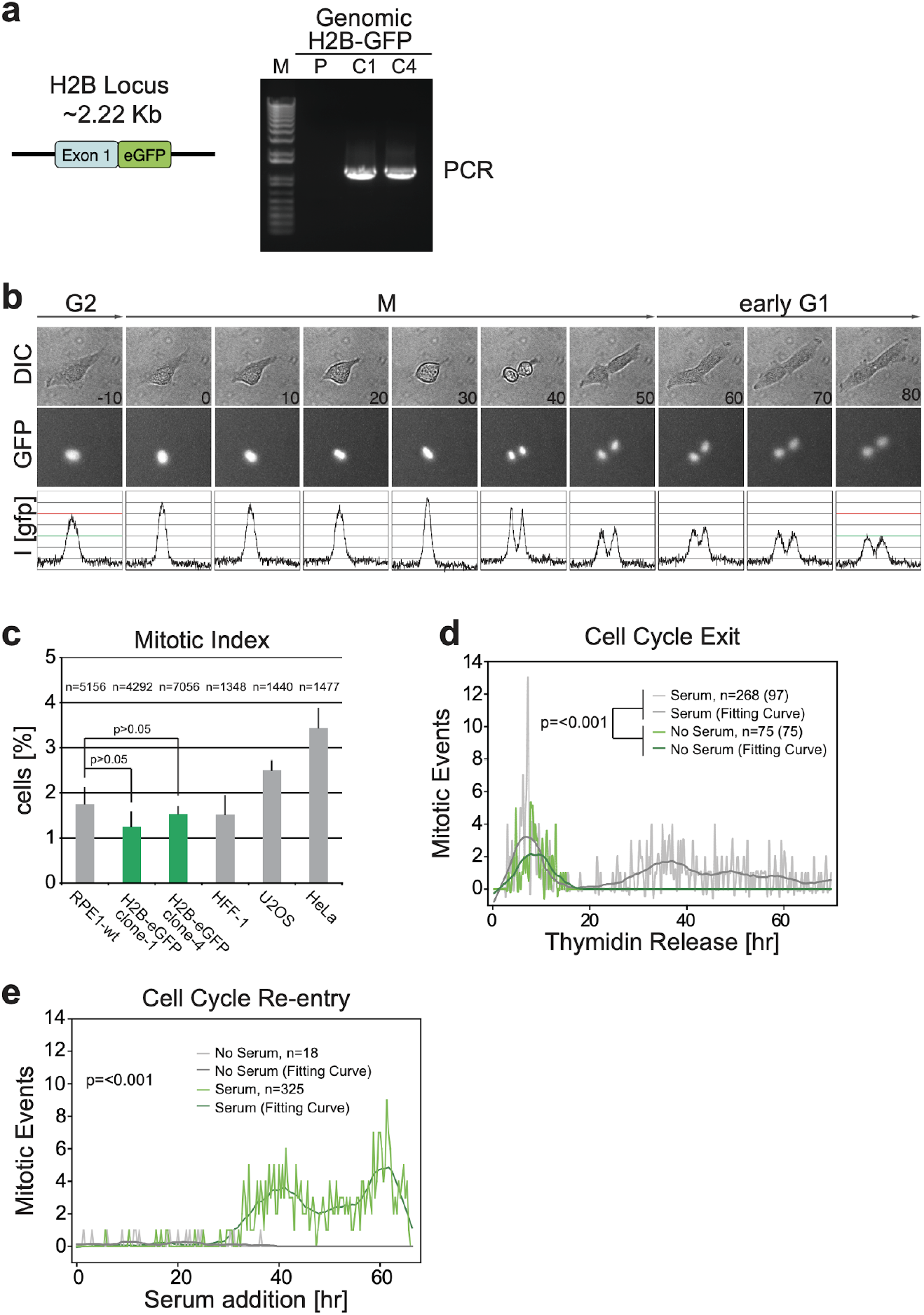
H2B-eGFP hTERT-RPE1 Cells. (**a**) H2B-GFP knock-in design, and site- and integration specific PCR to identify correctly recombined clones (C1, C4). (**b**) Time-lapse video microscopy of a mitotic H2B-eGFP cell. The G2 phase GFP intensity is double the intensity of the two daughter G1 cells (H2B is only transcribed and translated during S phase). (**c**) H2B-eGFP does not interfere with the cell cycle. The mitotic index is determined by video microscopy, comparing the parental line to two clones of H2B-eGFP, a primary human line (HFF), and two human cancer cell lines. (**d**) RPE1 H2B-eGFP cells were synchronized by Thymidine blockage and subsequently released into either full or serum-free media to follow cell cycle exit. (**e**) RPE1 H2B-eGFP cells were treated with serum-free media for 72 hr, then full media was added and cell cycle re-entry was followed by time-lapse video microscopy.

**Supplemental Figure S2. p27-mCherry hTERT-RPE1 Cells**.

(**a-c**) p27-mCherry/H2B-GFP hTERT-RPE1 cells were treated overnight under indicated conditions and assayed by Time-lapse video microscopy for both p27-mCherry (y axis) and H2B-GFP (x axis) levels. Five random images were taken (100 ms BF, 4,000 ms GFP, 10,000 ms mCherry) (**d**) Asynchronize p27-mCherry/H2B-GFP hTERT-RPE1 cells were assayed by Time-lapse video microscopy for both p27-mCherry (y axis) and H2B-GFP (x axis) levels using coordinates from a-c (above).

## Notes

### Competing Interest Statement

The authors have declared no competing interest.

